# Human papillomavirus 16 positive cervical cancer in Guatemala: The D2 and D3 sublineages differ in integration rate and age of diagnosis

**DOI:** 10.1101/2020.01.07.897546

**Authors:** Hong Lou, Joseph F. Boland, Edmundo Torres-Gonzalez, Anaseidy Albanez, Weiyin Zhou, Mia Steinberg, Lena Diaw, Jason Mitchell, David Roberson, Michael Cullen, Lisa Garland, Sara Bass, Robert Burk, Meredith Yeager, Nicolas Wentzensen, Mark Schiffman, Enrique Alvirez, Eduardo Gharzouzi, Lisa Mirabello, Michael Dean

**Author notes:** Contributed equally (Hong Lou, Joseph F. Boland, Edmundo Torres-Gonzalez, Anaseidy Albanez). Correspondence to Michael Dean, Ph.D., ATC, Room 134D, Laboratory of Translational Genomics, National Cancer Institute, Gaithersburg, MD 20877 USA.

## Abstract

Human papillomavirus (HPV) 16 displays substantial sequence variation; four HPV16 lineages (A, B, C, D) have been described, as well as multiple sub-lineages. To identify molecular events associated with HPV16 carcinogenesis we evaluated viral variation, the integration of HPV16, and somatic mutation in 96 cervical cancer samples from Guatemala. A total of 64% (60/94) of the samples had integrated HPV16 sequences, and integration was associated with an earlier age of diagnosis (P=0.0007) and pre-menopausal disease. HPV16 integration sites were broadly distributed in the genome but in one tumor, HPV16 integrated into the promoter of the interferon regulatory factor 4 (*IRF4*) gene, which plays an important role in the regulation of the interferon response to viral infection. The HPV16 D2 and D3 sub-lineages were found in 23% and 30% of the tumors, respectively and were significantly associated with adenocarcinoma. D2-positive tumors had a higher rate of integration (P=0.011), earlier age of diagnosis (P=0.012), and a lower rate of somatic mutation (P=0.03). Whereas D3-positive tumors are less likely to integrate, have later age-of-diagnosis, and a higher rate of somatic mutation. In conclusion, Guatemalan cervical tumors have a high frequency of the very high-risk HPV16 D2 and D3 sub-lineages and cervical cancer patients with these variants of HPV16 differ in histology, age of- diagnosis, integration, and somatic mutation frequency. In summary, related lineages of HPV16 have different features of oncogenicity.

## Introduction

High-risk human papillomavirus (HR-HPV) infection is responsible for nearly all cervical cancer (1). However, for unknown reasons, only <5% of these common HPV infections will progress to cervical precancer, which could lead to invasive cervical cancer (2). Other factors such as viral integration, viral variation, somatic mutations, genetic susceptibility, and host-virus interactions contribute to carcinogenesis (3–7). There are thirteen HPV types considered carcinogenic by IARC, with the most frequent being HPV16 (8). HPV16 is found in over 50% of the cervical cancers worldwide but displays substantial regional variation (9,10). HPV16 can be classified into four evolutionary-defined lineages (A, B, C, D), and 16 sublineages (A1-4, B1-4, C1-4, D1-4) (11), that vary in geographic distribution and disease risks (12–15). HPV16 sublineages differ by 0.5-1% of the viral genome, and the A4, D2 and D3 sublineages are associated with 9-28-fold higher risk of cancer and 10-130-fold risk of adenocarcinoma (15).

The oncogenicity of HPV is largely due to the E6 and E7 viral oncoproteins, essential to the life cycle of the virus (16). During an HPV infection, several transcripts are produced including an unspliced transcript producing E6 protein and an E6* spliced transcript generating the viral oncoprotein E7 (17,18). Recent studies show that HPV16 shifts to express predominantly the spliced, E7-producing transcript upon integration and that the E7 sequence is very highly conserved in cancers (3,7).

HPV DNA integration occurs in many sites in the human genome, with some preference for transcribed regions and common fragile sites (19–22). Integration into the genome of cancer cells can involve the deletion of all or portions of the viral E2 and E1 genes leading to deregulated expression of E6 and E7 viral oncogenes, which accelerates cell cycle entry, apoptosis, chromosome instability, and evasion of apoptosis (23). Inactivation of cellular tumor suppressor genes or activation of proto-oncogenes is rarely a direct consequence of HPV DNA integration (3,4,24–26); however, integration can induce chromosome instability (26,27). The rate of HPV integration is variable by HPV type, ranging from 55 to 95% of cervical tumors, with the highest rates observed for HPV18 and 45 (3,28). Integration is not required for HPV16 to cause invasive cancer, and episomal-only tumors are found as well as tumors with both integrated and episomal HPV16 sequences (29).

Our previous study demonstrated that up to 33% of cervical cancers from Latin America carry *PIK3CA* mutations, mainly at E542K and E545K, two specific hot spots in the helical domain of the *PIK3CA* gene (30). These mutations are part of a spectrum of mutations caused by the APOBEC family of enzymes activated by viral infection (31,32). Also, there are at least 14 cellular genes frequently mutated in cervical tumors, including *EP300*, *PTEN*, *HLA-B*, *KRAS*, and *FBXW7* (3,4,30,31,33).

The goal of this study was to further dissect the molecular events involved in cervical cancer in HPV16-positive tumors. Using whole virus sequencing, viral DNA capture’ and integration analysis, and somatic sequencing we have characterized the integration status of the major HPV16 sublineages.

## Materials and Methods

### Sample Collection

Ninety-six HPV16-positive samples from Guatemala were selected from a total of 298 cervical tumors collected between 2011 and 2013 at the Instituto de Cancerologia (INCAN), the central reference hospital for adult cancer therapy in the country. Selected samples were confirmed HPV16 positive with at least 1 ug of tumor DNA for sequence capture. All patients, between 25- and 85-year-old, presenting for confirmation of the diagnosis of cervical cancer were invited to participate. Written informed consent was obtained from the patients. Protocols were approved by the Ethical Review Committees of the local hospitals in Guatemala City, Guatemala, and the Office of Human Research Subjects, NIH, in accordance with the Belmont Report. Cervical tumor biopsy tissues were immediately stored in RNAlater (QIAGEN) at −20°C until extraction; blood samples were stored at −20°C and a questionnaire on reproductive history, smoking, oral contraceptive use, and family cancer history was administered.

### DNA and RNA Extraction, and HPV Genotyping

DNA and RNA were extracted from the cervical cancer tissues (5-10 mg) using the AllPrep DNA/RNA Micro Kit (QIAGEN) as described by the manufacturer. HPV type was determined by touch-down PCR using degenerate Broad-Spectrum (BS) GP5+ and GP6+ primer amplification and Sanger sequencing (Schmitt M. et al. J Clin Microbiol. 2008; 46(3):1050–9). The detailed method for the determination of HPV genotyping is described (30).

### Quantitative RT-real-time PCR

One microgram of total RNA was reverse transcribed into cDNA using the Transcriptor First Strand cDNA Synthesis Kit (Roche) with the oligonucleotide (dT)_18_ primer according to the manufacturer’s instructions (30,34). Real-time PCR was performed for E6 and E7 HPV transcripts, using gene and type-specific primers, in the presence of SYBR green. Relative expression levels of E6 and E7 were normalized using the *β-actin* gene as described (35). The E6 unspliced/spliced ratio was determined using the following formula: E6 mRNA value (unspliced) divided by (E6*/E7 mRNA value minus E6 mRNA value).

### Determination of *PIK3CA* and other somatic mutations

Somatic mutations were identified using exome and ultra-deep targeted sequencing as described in (30).

### HPV DNA Capture using the NimbleGen Array

DNA preparation: 200 ng genomic DNA was purified using Agencourt AMPure XP Reagent (Beckman Coulter, Inc.) according to the manufacturer’s protocol. HPV library preparation: full-length HPV genomes were amplified pre-hybridization by ligation-mediated PCR consisting of one reaction containing 20 μl library DNA, 25 μl 2x KAPA HiFi HotStart ReadyMix, and 5 μl 10x Library Amplification Primer Mix. PCR was subjected to the following cycling conditions: 98°C for 45 s, followed by 5 cycles of 98°C for 15 s, 60°C for 30s, 72°C for 30 s. The last step was an extension at 72°C for 1 minute. The amplicons were purified by Agencourt AMPure XP Reagent (Beckman Coulter, Inc.) according to the KAPA provided protocol. Liquid Phase Sequence Capture: HPV DNA capture was performed with NimbleGen’s SeqCap EZ Custom Library containing probes for all HR HPV types (Roche NimbleGen, Inc.,) according to the manufacturer’s protocol. The DNA mixture was hybridized at 47°C for 64 to 72 hours and washing and recovery of captured DNA were performed as described in NimbleGen SeqCap EZ Library SR Protocol. The captured DNA was amplified by ligation mediated PCR (Beckman Coulter, Inc.) according to NimbleGen SeqCap EZ Library SR Protocol. The captured DNA was quantified via Kapa’s Library Quantification Kit for Ion Torrent (Kapa Biosystems) on the LightCycler 480 (Roche, Indianapolis, IN, USA). All sequencing was performed using an Ion Torrent Proton Sequencer using. The sequence captured both episomal and integrated HPV16 DNA. The integrated reads map to both human sequence and HPV16 sequence and determine the breakpoint for HPV integration.

### HPV16 genome sequencing and sub-lineage determination

We used a custom Thermo Fisher Ion Torrent AmpliSeq HPV16 panel approach to amplify the entire 7906 bp HPV16 genome as previously described (36). HPV16 variant lineage assignment was based on the maximum likelihood tree topology constructed in MEGA, including 16 HPV16 variant sublineage reference sequences (10), and lineage assignments were confirmed with SNP patterns. The detailed methodology was described previously (15).

### Confirmation of IRF4 integration by PCR, cloning, and sequencing

HPV16 integration into the *IRF4* gene was confirmed by PCR using GoTaq Long PCR Master Mix kit (Promega U.S.), with 5’-AAGCATGTCAGACACGCAGA- 3’as a forward primer, and 5’- GGAAGACGGAGGAATGGTCC- 3’as a reverse primer. DNA samples from blood and tumor tissues were assessed; normal donor blood DNA was used as a negative control (Figure S1A).

### Statistical Analysis

Unpaired T-test, Pearson’s chi-squared test, and Fisher’s exact test statistical analyses were performed using GraphPad Prism version 7 for Windows; P<0.05 was regarded as significant.

## Results

### Analysis of HPV16 sequence and integration

To determine the genomic characteristics of HPV16 positive tumor samples from an understudied population, we selected 94 HPV positive tumor samples from 298 subjects in an ongoing hospital-based case study of advanced cervical cancer (94% stage IIB or higher) in Guatemala. The selection criteria were HPV16 positive by amplification and sequencing of a region of the L1 gene, and adequate DNA (1 ug) for sequence capture. Both squamous cell carcinoma (SCC) and adenocarcinoma cases were included. HPV DNA was captured and reads mapped to both the human and HPV16 genome and overlapping amplicon sequencing of the virus was also performed (30,36). Complete HPV16 sequences were aligned, and phylogenetic analyses were used to determine HPV16 sublineages (15). HPV integration sites were determined from HPV-human junction reads based on captured HPV DNA and targeted human gene sequencing was previously reported (30). Samples with high E6 and E7 coverage and low coverage in multiple E2 and E1 gene amplicons in the amplicon sequencing were determined to have integration without episomal DNA based on HPV16 genome capture.

A total of 64% (60/94) of the tumors had integrated HPV16 with equal numbers having both episomal DNA and integration (30/60) and only integrated HPV16 (30/60). (**Figure 1**). There was a significant decrease in the frequency of integration (82% to 48%, P=0.03) with increasing age of diagnosis and with post-menopausal disease (P=0.009) (**Table 1**). The median age at diagnosis for patients with tumors with HPV16 integration was younger (median=48, SD=12) compared to patients with tumors without integration (median=57, SD=13) (P=0.002, Mann-Whitney test) (**Figure 2A**). Integration was not significantly correlated with histological subtype (adenocarcinoma or squamous cell carcinoma), the number of pregnancies, smoking, cooking method, oral contraceptive use, or age at menarche (**Figure 1, Table 1**).

**Figure 1.**
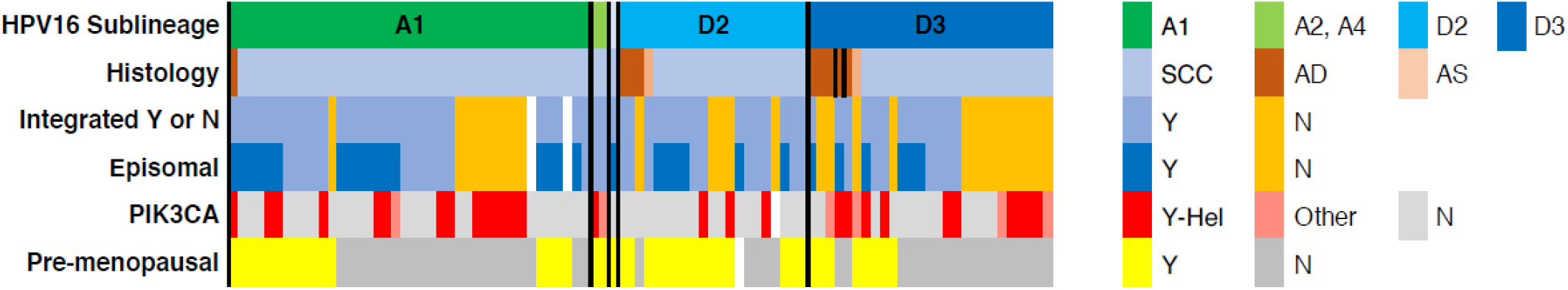
The landscape of HPV16 Cervical Tumors in Guatemala. Characteristics of HPV16 tumors are shown including HPV16 sublineage, histology, menopausal status, HPV integration status and PIK3CA mutation status. SCC, squamous cell carcinoma; AC, adenocarcinoma, ASC, adenosquamous; Pre, pre-menopausal; Men, post-menopausal; Int, HPV integrated only; Int/Ep, integrated and episomal; Ep, Episomal virus only; Mut-H, *PIK3CA* helical domain mutation; Mut, any *PIK3CA* mutation; WT, wild type for *PIK3CA*.

**Table 1.**
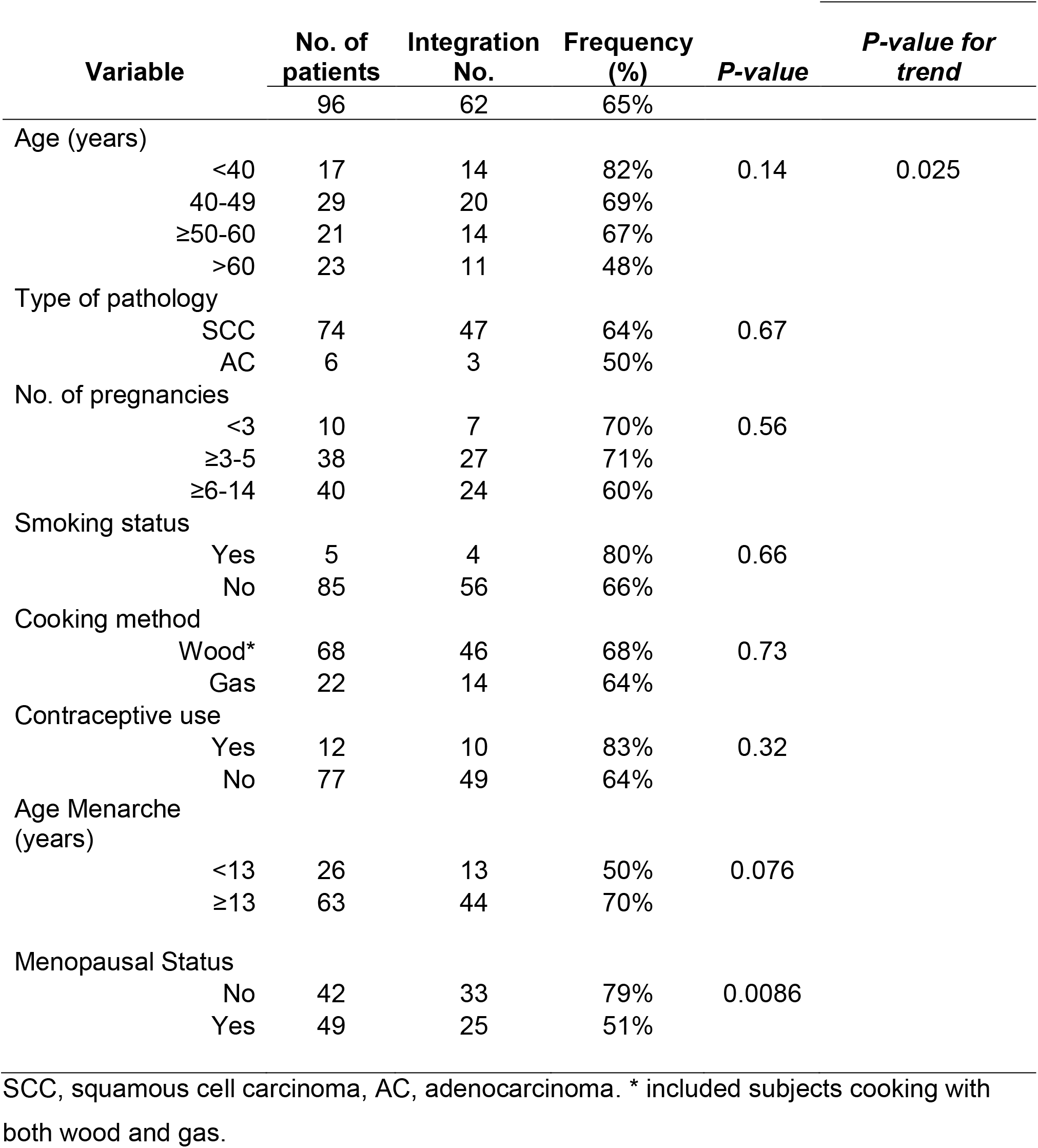
Demographic and clinicopathologic characteristics of cervical cancer and HPV16 integration.

**Figure 2.**
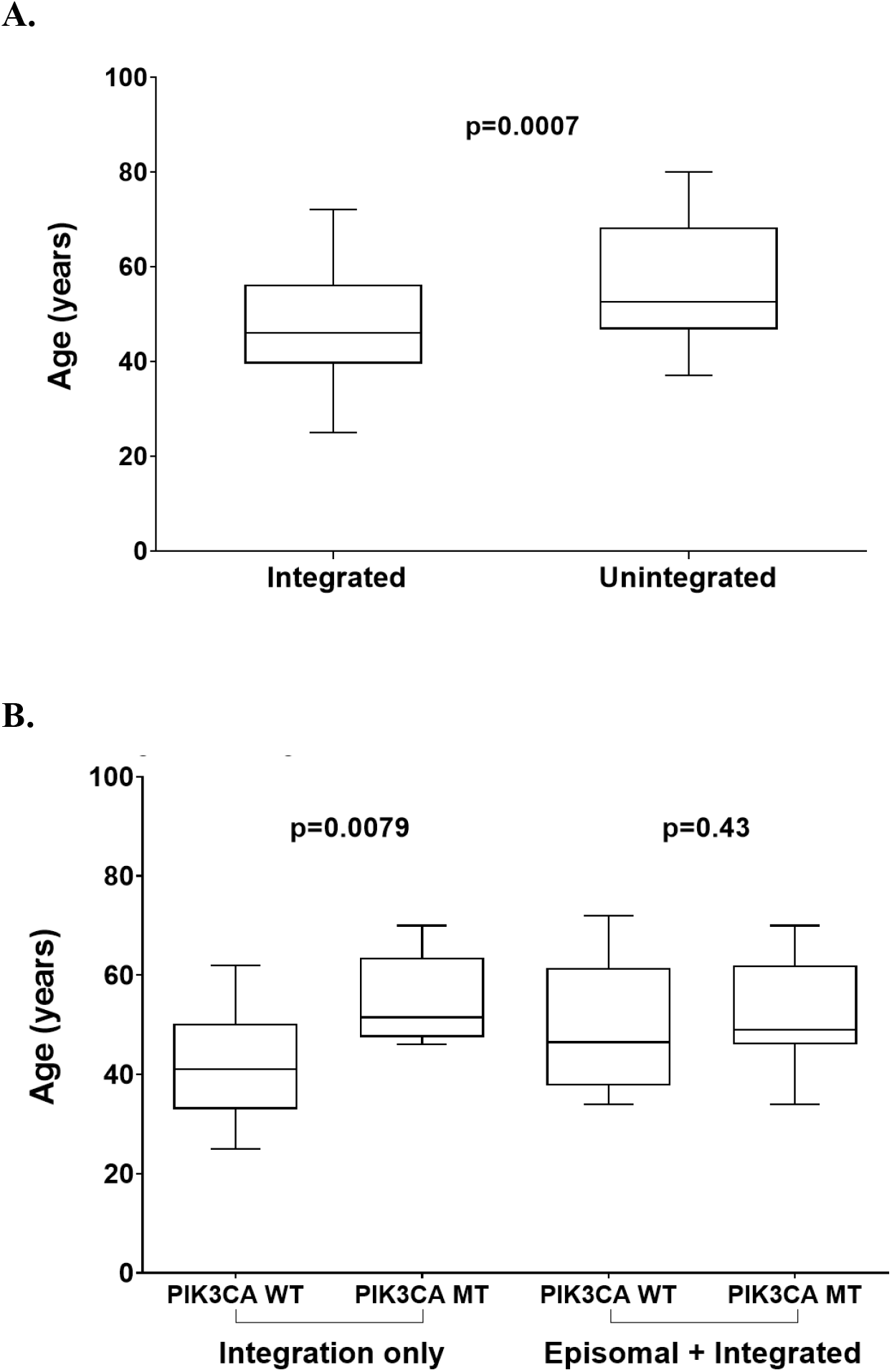
HPV integration and age of diagnosis. A, the age of diagnosis of patients with integrated HPV (includes integrated only and integrated and episomal) is compared against unintegrated samples. B, the age of diagnosis of integration only and episomal in integrated samples is compared by *PIK3CA* mutation status.

### Location of HPV16 integration sites

To confirm that there were not specific or recurrent sites of HPV16 integration, we mapped integration location. In the 60 patients with integration, a dispersed distribution of sites in the human genome was observed with sequences localized to promoter, intron, exon, and intergenic regions with no two integrations in the same gene (**Table S1, S2**). Integration in genes not previously reported includes the promoter of *EGR3*, *IRF4*, and *PLA2G4E*, and exons of *LOC100506022*, *FOXK1*, *GAPDH*, and *TBC1D24* (**Table S3**). The *IRF4* integration was confirmed using PCR, cloning, and sequencing (**Figure S1A, S1B**). Integration into promoter regions occurred predominantly in D2 or D3 sublineage-containing tumors and exon regions for A1-positive tumors, but this was not significant (**Table S3**).

### Somatic mutations

In HPV integrated samples, there were fewer mutations in *PIK3CA* (29% in integrated vs. 53% in unintegrated tumors, P=0.02). This was also true for mutations in known cervical cancer driver genes including *PTEN*, *TP53*, *STK11 HRAS*, *MAPK1*, and *FBXW7* (45% vs. 68%, P=0.03). There was also a correlation between age of diagnosis and somatic mutation status, with a higher frequency of *PIK3CA* mutations in older onset patients (**Figure 2B**).

### HPV16 Sub-lineage frequency and variants

To understand the role of viral variation we analyzed HPV16 sublineages and clinical and molecular characteristics of the patients. The sub-lineage A1 (44%, 40/91) was the most frequent followed by D3 (30%, 27/91), D2 (23%, 21/91) and only infrequent numbers of A2, A4, and D1 were observed (**Table 2**). Adenocarcinomas had almost exclusively A4, D2 and D3 viruses (11/12, 92%), whereas only 37/77 SCC tumors (48%) were positive for A4, D2, or D3 (P=0.002) (**Table 2**).

**Table 2.** Frequency of HPV16 sub-lineages and histologic types.

| Sublineage | Total No.<br>(n=88) | Frequency | Squamous<br>cell<br>carcinoma | Adenocarcinoma | Freq.<br>Adenocarcinoma |
| --- | --- | --- | --- | --- | --- |
| A1 | 40 | 45% | 39 | 1 | 3% |
| D2 | 21 | 24% | 17 | 4 | 19% |
| D3 | 27 | 31% | 19 | 7 | 27% |
|  | 88 |  | 75 | 12 |  |

Not shown are single samples of A2, A4, and D1 all squamous cell carcinoma.
Adenocarcinoma includes adenosquamous carcinoma.

Analysis of rare variants in HPV16 sequences identified only 1 variant (F57V) in the E7 gene in 91 samples as compared to 9 variants (Q10E, H31Y, H31N, I34R, E36Q, R55W, S78C, C118R, and T156A) in the E6 gene (**Table S4**). This is consistent with hypovariation in the E7 gene in cancers described in our previous study (7).

To identify differences associated with the more common HPV16 A1, D2, and D3 sublineages, we analyzed the data by age-of-diagnosis, integration frequency, and presence of a *PIK3CA* or other somatic mutation. Patients with the D2 and A1 sublineages had a significantly higher integration frequency (76%, 16/21 and 78%, 31/40, respectively) than those with D3 (44%, 12/27; P=0.01) (**Figure 3A**). Also, patients with D2 had a significantly younger age of diagnosis than A1 and D3-containing patients and are more often pre-menopausal (P=0.017) (**Figure 3B**). D2-containing tumors had a significantly lower rate of somatic mutation in *PIK3CA* than D3 (P=0.007) (**Table 3**), however, this was not significant for all somatic driver mutations (data not shown).

**Table 3.**
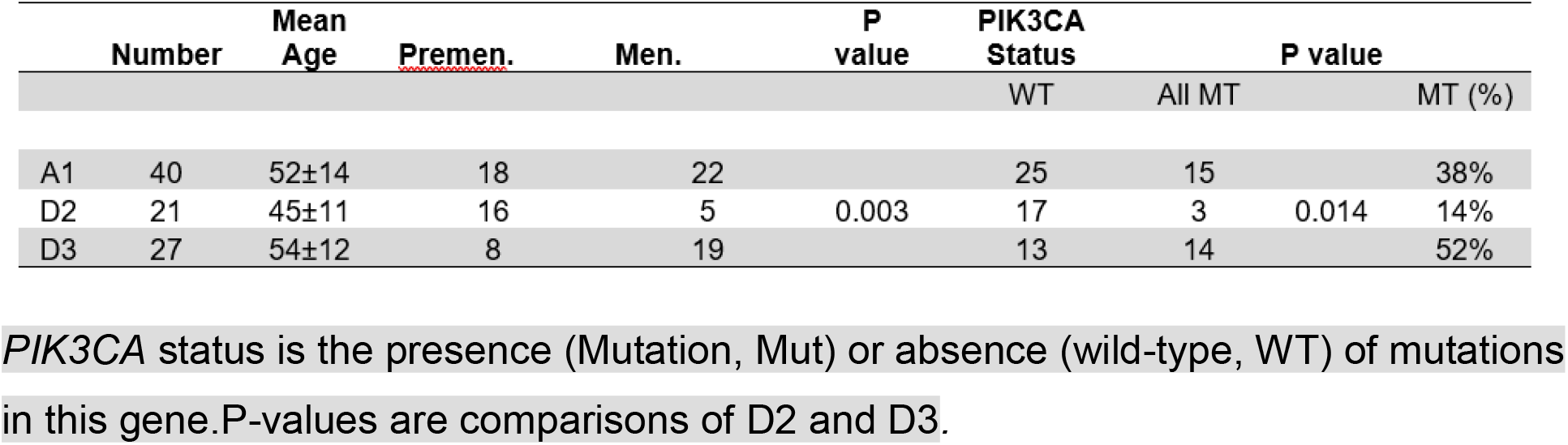
Correlation between HPV16 sublineage, age, and *PIK3CA* mutation.

**Figure 3.**
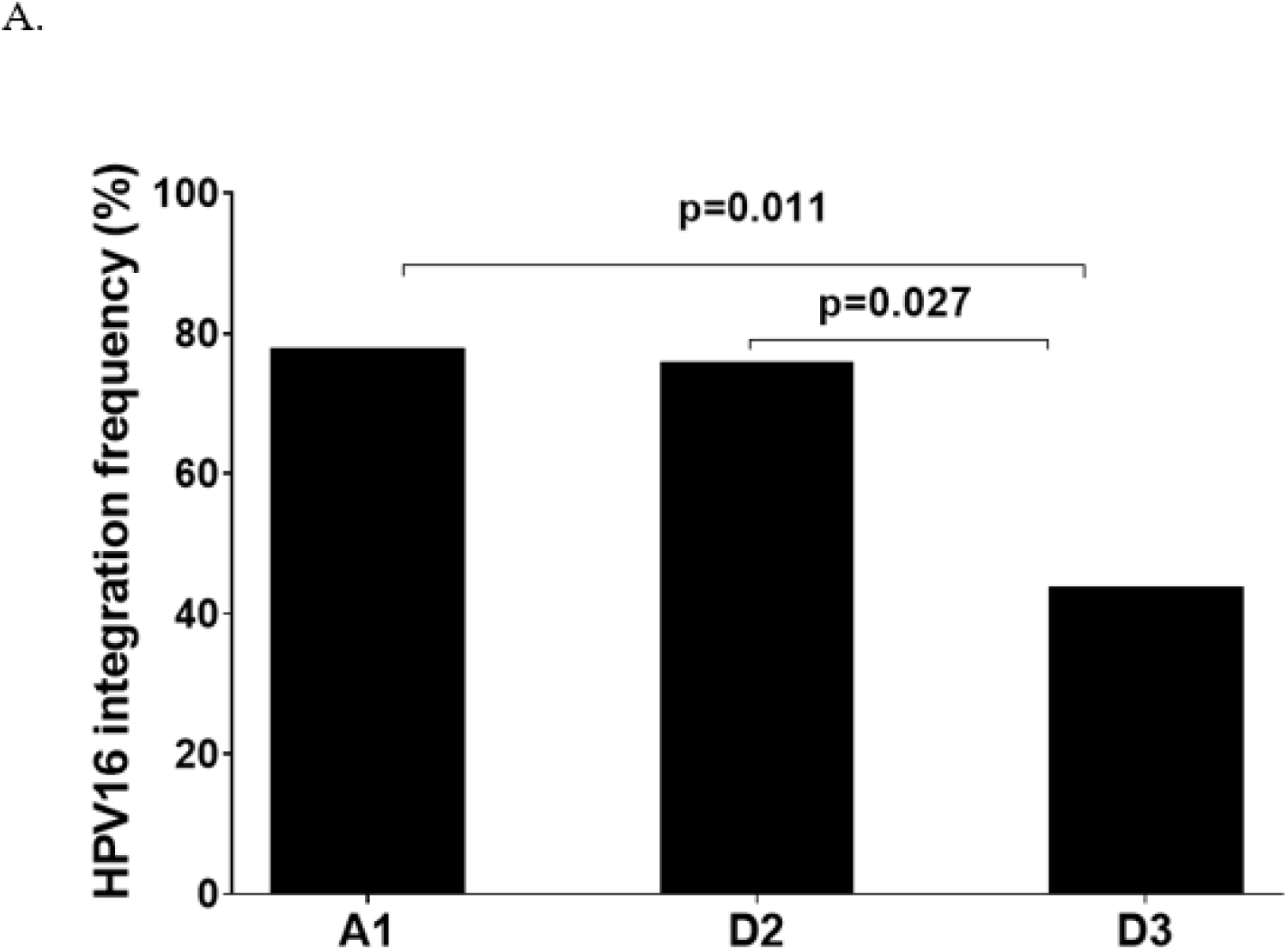
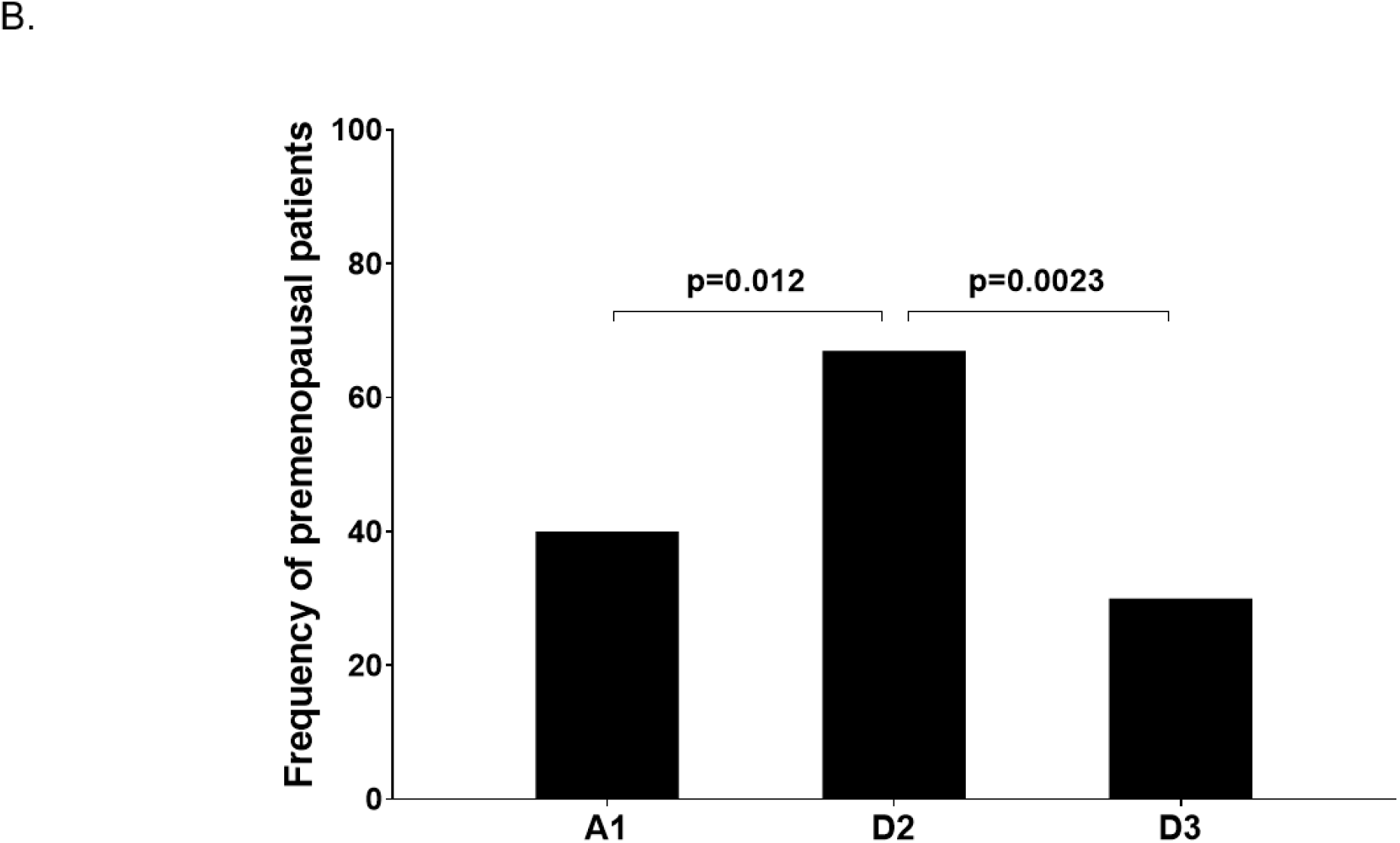
HPV16 sublineage, integration and menopausal status. A, the frequency of integration is displayed by HPV16 sublineage. B, The frequency of pre-menopausal diagnosis is displayed by the HPV16 sublineage.

Because we observed significant differences between the D2 and D3 sublineages, we examined the lineage-defining amino acid changes between these sublineages. There are 11 amino acid differences between D2 and D3 as compared with the reference A1. These alterations involve 5 proteins, E1, E2, E5, L1, and L2, but 5/11 (45%) alterations are in the E2 gene (**Table S5**). The A1, D2 and D3 E2 proteins show five amino acid positions that are altered at residues 136, 142, 157, 211 and 221 (**Table S6**).

## Discussion

To comprehensively study integration events in HPV16 positive cervical tumors, we used HPV DNA capture and HPV whole-genome sequencing on 96 advanced-stage cervical tumors from Guatemala. We analyzed the HPV16 sequence, sublineage distribution, and viral integration frequency and sites of viral insertion. The integration frequency we observed, 65% of tumors, is consistent with other reports (24,28,37), and HPV16 integration was associated with a significantly younger age of diagnosis, as seen previously (28,38). The location of integrated HPV16 was dispersed throughout the human genome, but in selected tumors may disrupt potential cancer-related genes such as *FHIT*, *GLI2*, and *IRF4*. Our sample has a significant fraction of D2 and D3 sublineage-containing tumors allowing us to distinguish differences in age of diagnosis, integration and somatic mutation in these two highly carcinogenic forms of HPV16.

Recent progress in determining HPV whole-genome sequences at high-throughput from clinical samples have revealed extensive diversity in HPV16 genome sequences and allowed for comprehensive classification of HPV16 variants and sublineages (10,36). The A4, D2, and D3 sublineages have been found to have a higher risk for cancer, particularly for adenocarcinoma (15). There were only 8 D2 and 4 D3 HPV16 samples in the TCGA study, providing little information on the molecular details of these cancers. Our study provides information on age-of-diagnosis, histology and somatic mutations for 21 additional D2 and 27 D3-containing tumors. This sample size, although modest, was sufficient to determine that D2 and D3-tumors are distinct regarding integration (lower in D3), age-of- diagnosis (earlier in D2) and percentage of *PIK3CA* mutations (higher in D3). Combining our data with two other studies shows that adenocarcinoma is consistently higher in A4, D2, and D3 compared to A1 (**Figure 4A**). Also, the integration rate was lower in HPV16 A1 and HPV31, intermediate in HPV16 D2 and D3 and high in HPV18 and HPV45 when surveying other reports (**Figure 4B**).

**Figure 4.**
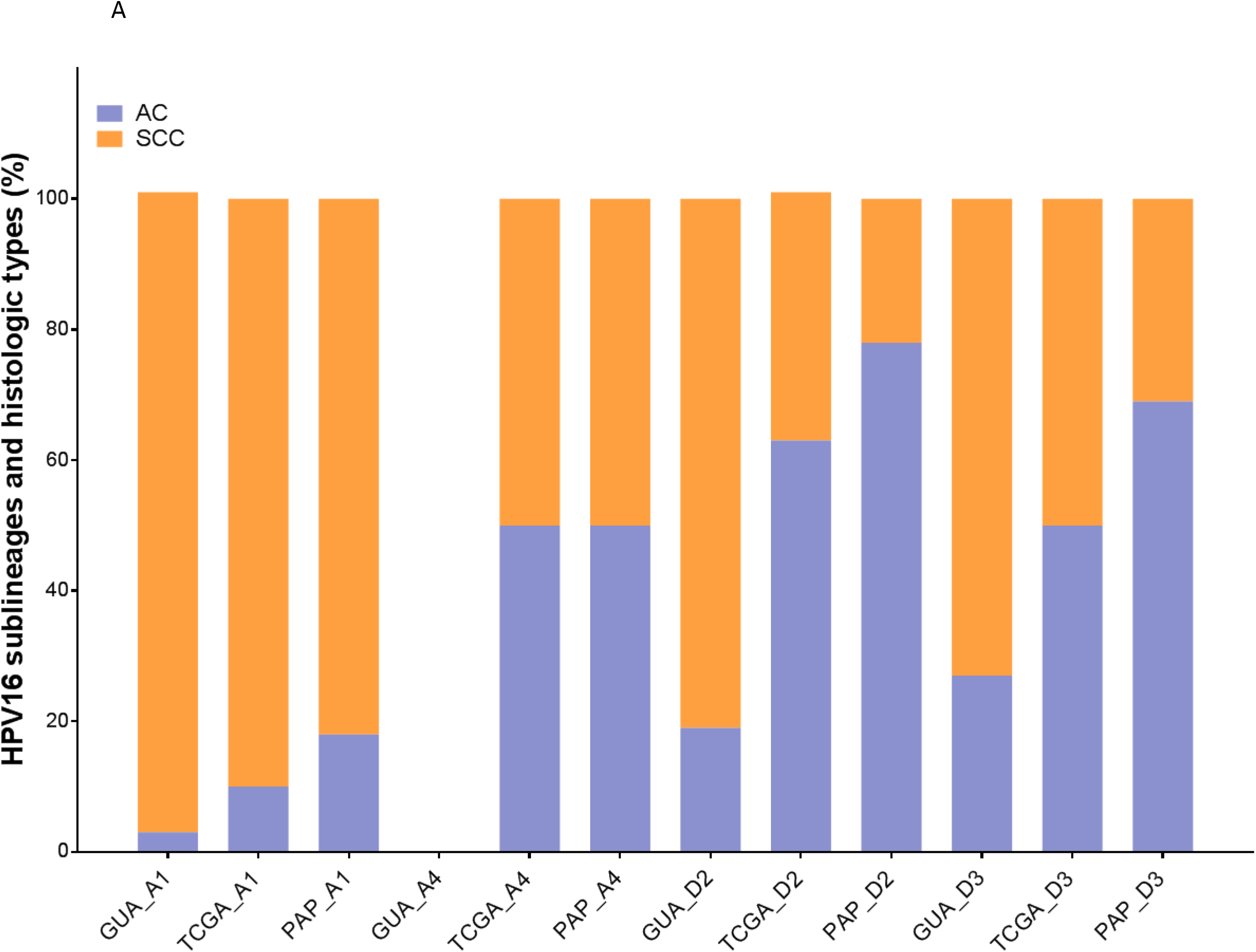
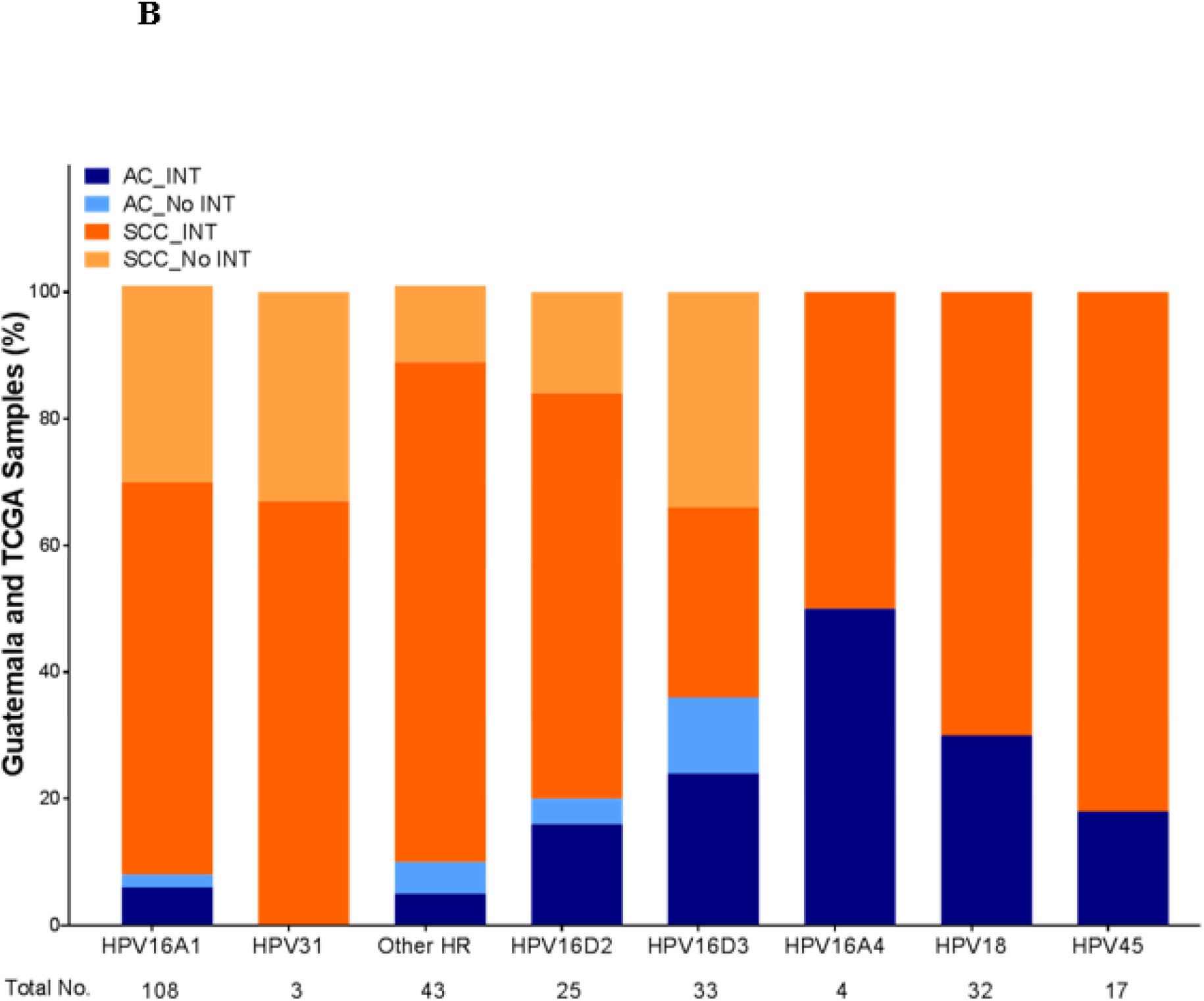
Comparison of adenocarcinoma percentage in Guatemala and other studies. A, The percentage of adenocarcinoma (AC) versus squamous cell carcinoma (SCC) is shown by HPV16 sublineage in the current study compared to that of TCGA (3) or the PAP cohort (15). B, combined integration and histology status of US and Guatemalan tumors for HPV16 sublineages, HPV18, HPV31, and HPV45.

The biological basis for the dramatic difference in carcinogenicity, especially for adenocarcinoma of the D2 and D3 sublineages, compared to A1 is important to understand. While there are no differences in E7 between A1 and D2 or D3, there are 3 amino acid differences in the E6 gene and functional studies indicate that these changes increase transforming ability (39). There are also a considerable number of amino acid differences (3–16) between A1 and D2 and D3 in other viral proteins, with the most variable E1, E2, and L1. Because the early genes E2, E1, E5 are required for viral replication, transcription, cellular transformation, initiation of neoplasia and maintenance of the viral genome in infected cells, they represent the best candidates for the difference in carcinogenicity. Nevertheless, because of the high correlation of changes throughout the genome, it is hard to infer causality of specific mutations and/or genes.

Our finding of substantial differences in the age of diagnosis, integration and somatic mutation between D2 and D3 tumors indicates that a careful comparison of these two sublineages would be informative. Almost half of the amino acid alterations between D2 and D3. are in the E2 gene, a transcription factor that binds to the viral origin of replication and the major promoter for E6 and E7 oncogene expression. These six amino acid changes at five sites are in the E2 transactivation domain (H136Y, E142D, L157M, and L157I) and the hinge domain (I211T, A221T). We propose that the E2 gene is a candidate for a gene that contributes to the higher rate of cervical cancer and adenocarcinoma seen with HPV16 D2 and D3, but direct functional analyses would be required to demonstrate this and non-coding variants are also likely to contribute to functional differences (7).

## Conclusions

In conclusion, we have established that a high proportion of HPV16 isolates in Guatemala are sub-lineage D2 and D3, known to have high cancer and adenocarcinoma risk. D2-positive patients have a significantly higher integration frequency, significantly younger age and significantly lower *PIK3CA* mutation frequency than D3. Further studies of these isolates and tumors may yield information on the mechanism of adenocarcinoma formation and the role of integration in cervical carcinogenesis.

## Acknowledgments

We thank the patients that agree to participate in this study and the physicians and staff that helped recruit subjects and collect samples. We thank Esther Avila, Lineth Boror, and Paty Zaid for sample and data collection and shipping. Supported in part by the NIH Intramural Research Program.

## Contributions

MD and LM contributed to study conception and design, WZ, MS, JM, DM, SB, and JB to development of methodology, AA, EA, and EG to acquisition of data and samples, MD, LM, and ET-G to analysis and interpretation of data, HL, RB, MW, NW, MS, LM, and MD to writing, review or revision of the manuscript.

## Notes

The authors have no conflicts of interest to report.

