## Supplemental figures_Tables for "Human papillomavirus 16 positive cervical cancer in Guatemala: The D2 and D3 sublineages differ in integration rate and age of diagnosis"

**Supplemental Tables and Figures**

**Table S1. Sites of HPV16 integration**.

| **Integration in class** | **Number of events** | **Chromosome** |
| --- | --- | --- |
| **Promoter** | 4 | Chr. 6, 8, 15, 17 |
| **Exon** | 5 | Chr. 2, 6, 7-9 |
| **Intron** | 42 | Chr. 1-4, 6, 8, 9, 10, 11, 12, 13, 14, 15, 16, 17, 18, 19, 20, 21 |
| **intergenic** | 21 | Chr. 1-4, 6, 8, 10, 11, 14, 15, 21 |

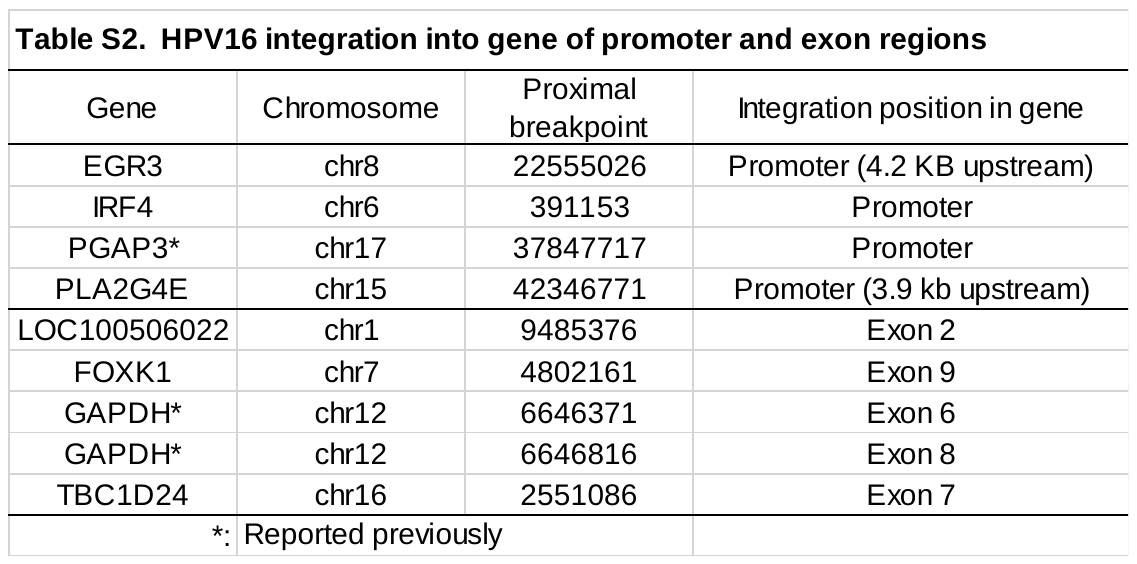

| **Table S3. Correlation between HPV16 integration site and clinical data** | | | | | |  |
| --- | --- | --- | --- | --- | --- | --- |
| Gene | Integration position | Integration status | HPV16 sublineage | Age | Age at Menarche | Age at first Pregnancy |
| IRF4 | promoter | Ep and Int* | A1 | 36 | 14 | 19 |
| EGR3 | promoter | Ep and Int | D3 | 55 | 15 | 16 |
| PGAP3 | promoter | Ep and Int | D3 | 46 | 11 | 23 |
| PLA2G4E | promoter | Ep and Int | D2 | 48 | 10 | 19 |
| FOXK1 | Exon | Int only | D1 | 36 | 17 | 25 |
| LOC100506022 | Exon | Int only | A1 | 60 | 15 | 20 |
| GAPDH | Exon | Int only | A1 | 46 | 12 | 20 |
| TBC1D24 | Exon | Int only | A1 | 25 | 13 | NA |
| * | Episomal (Ep) and Integrated (Int), NA=not applicable | | |  |  |  |

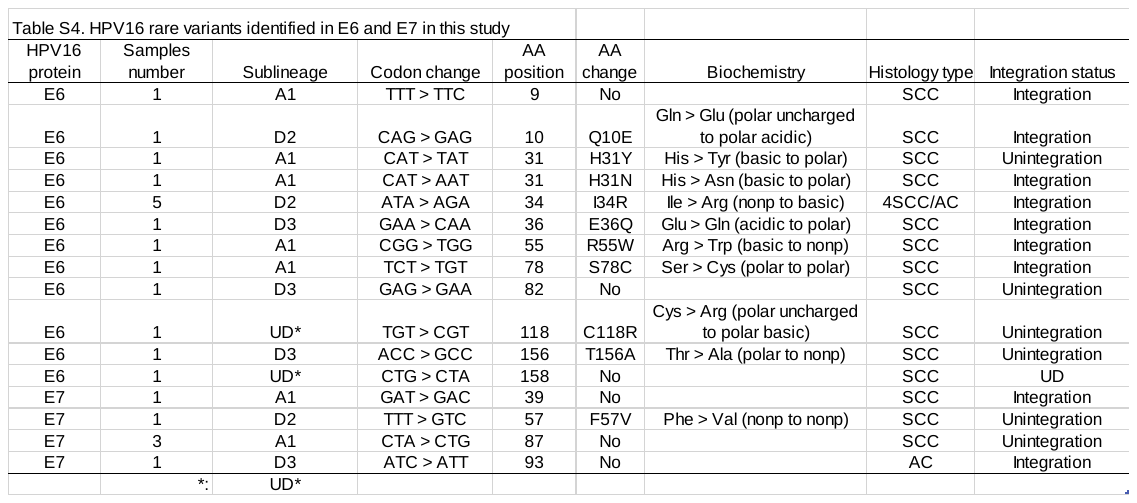

UD=undetermined

| **HPV16 Sub-lineage** | **Position of AA change** | **Reference AA (A1)** | **AA change** | **Variant** | **HPV16 protein** | **Protein domain** |
| --- | --- | --- | --- | --- | --- | --- |
| D2 | 333 | Y | C | Y333C | E1 | DBD |
| D2 | 136 | H | Y | H136Y | E2 | Transactivation |
| D3 | 142 | E | D | E142D | E2 | Transactivation |
| D2 | 157 | L | M | L157M | E2 | Transactivation |
| D3 | 157 | L | I | L157I | E2 | Transactivation |
| D3 | 211 | I | T | I211T | E2 | Hinge Domain |
| D2 | 221 | A | T | A221T | E2 | Hinge Domain |
| D2 | 40 | T | A | T40A | E5 | TMD2 |
| D3 | 207 | N | T | N207T | L1 |  |
| D3 | 415 | T | S | T415S | L1 |  |
| D2 | 351 | T | P | T351P | L2 |  |

**Table S5. Amino acid differences from the reference HPV16 A1 sublineage.**

DBD, DNA-binding domain, TMD2, transmembrane domain 2.

**Table S6. E2 gene haplotypes for HPV16 A1, D2, and D3 sub-lineages**

|  | **AA position** | |  |  |  |  |  |  |  |  |  |  |  |  |  |  |  |  |
| --- | --- | --- | --- | --- | --- | --- | --- | --- | --- | --- | --- | --- | --- | --- | --- | --- | --- | --- |
|  | **35** | **135** | **136** | **142** | **143** | **157** | **165** | **203** | **208** | **211** | **219** | **221** | **232** | **254** | **271** | **310** | **341** | **344** |
| A1 | H | T | H | E | A | L | R | N | P | I | P | A | E | T | F | T | W | D |
| D2 | Q | K | Y | . | T | M | Q | D | A | . | S | T | K | N | V | K | C | E |
| D3 | Q | K | . | D | T | I | Q | D | A | T | S | . | K | N | V | K | C | E |
| Domain |  |  | TA | TA |  | TA |  |  |  | Hinge |  | Hinge |  |  |  |  |  |  |

TA, Transactivation domain

**Supplemental Table S7. Frequency of HPV sublineages and histological types**

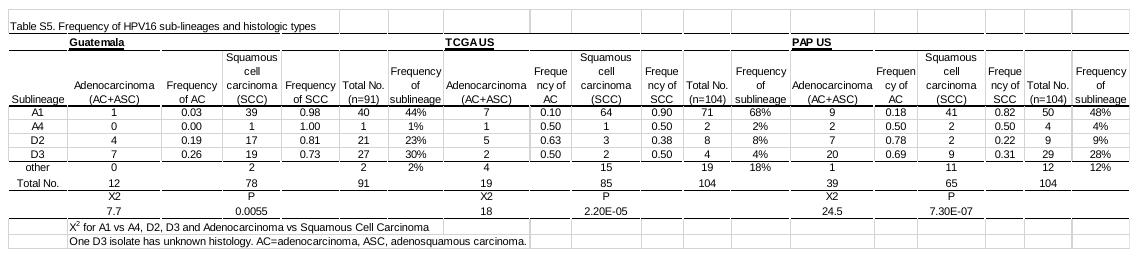

**Supplemental Figure S1A**

C Bl Tum

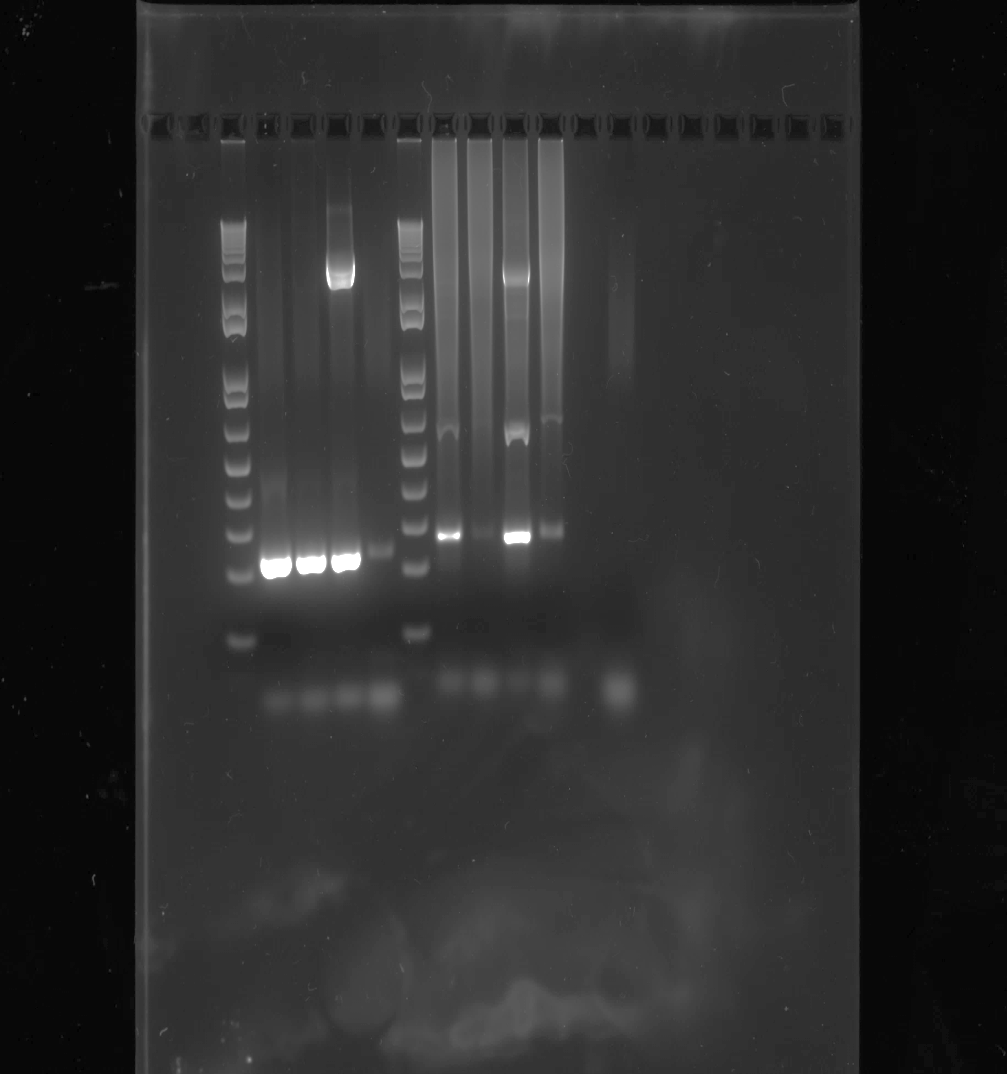

Normal Donor

BJ200262 Blood

BJ200262 CC Tissue

2,000bp

200bp

HPV16 integrated IRF4

IRF4

Chr6:391153

HPV (E5+L2)

Chr6:391169

Forward primer

Reverse primer

TSS

**IRF4**

**Supplemental Figure S1B**

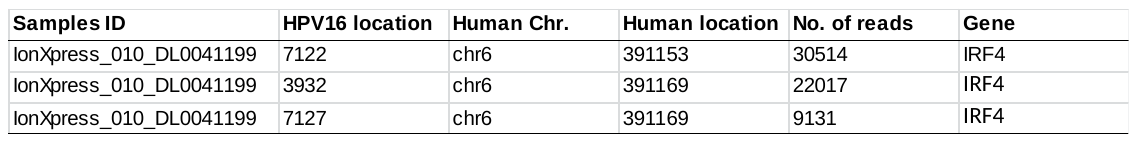

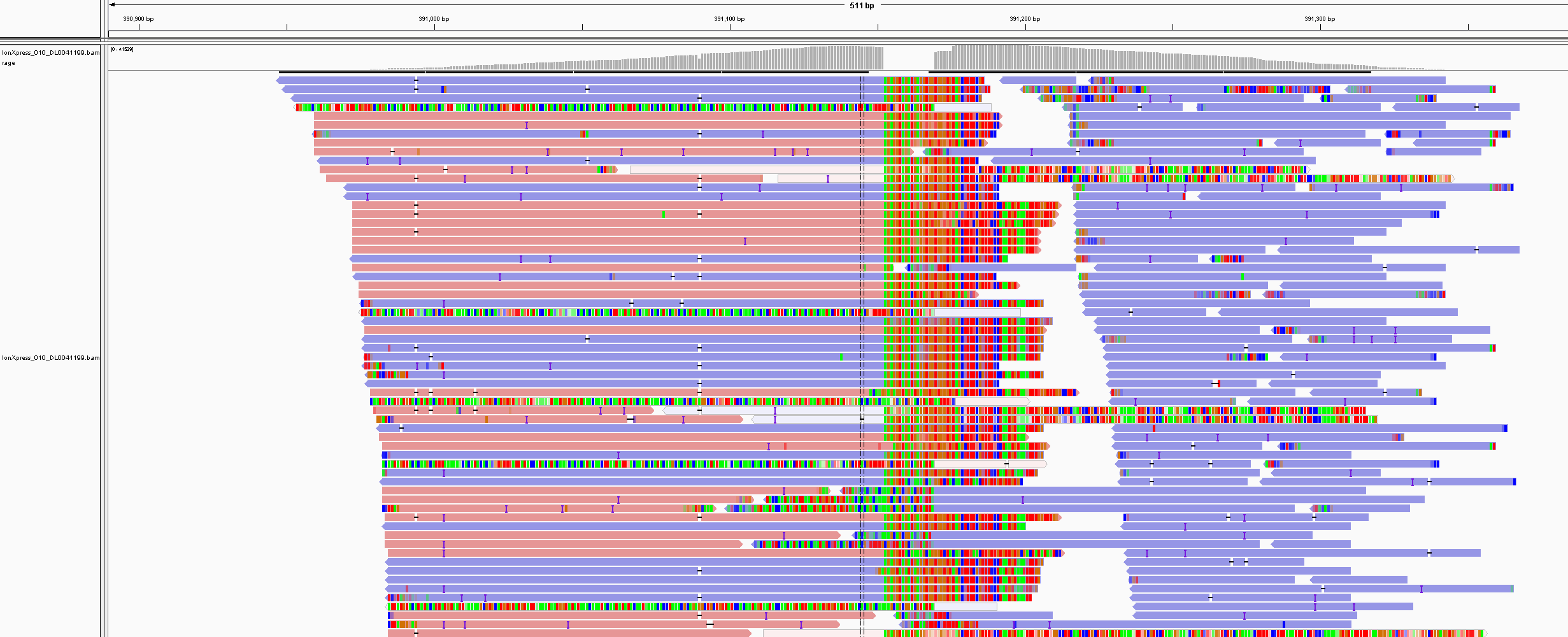
